# Characterizing nanometric thin films with far-field light

**DOI:** 10.1101/2022.08.15.503956

**Authors:** Hodaya Klimovsky, Omer Shavit, Carine Julien, Ilya Olevsko, Mohamed Hamode, Yossi Abulafia, Hervé Suaudeau, Vincent Armand, Martin Oheim, Adi Salomon

## Abstract

Ultra-thin, transparent films are being used as protective layers on semiconductors, solar cells, as well as for nano-composite materials and optical coatings. Nano-sensors, photonic devices and calibration tools for axial super-resolution microscopies, all rely on the controlled fabrication and analysis of ultra-thin layers. Here, we describe a simple, non-invasive, optical technique for simultaneously characterizing the refractive index, thickness, and homogeneity of nanometric transparent films. In our case, these layers are made of the biomimetic polymer, My-133-MC, having a refractive index of 1.33, so as to approach the cytosol for biological applications. Our technique is based on the detection in the far field and the analysis of supercritical angle fluorescence (SAF), i.e., near-field emission from molecular dipoles located very close to the dielectric interface. SAF emanates from a 5-nm J-aggregate emitter layer deposited on and in contact with the inspected polymer film. Our results compare favorably to that obtained through a combination of atomic force and electron microscopy, surface-plasmon resonance spectroscopy and ellipsometry. We illustrate the value of the approach in two applications, (*i*), the measurement of axial fluorophore distance in a total internal reflection fluorescence geometry; and, (*ii*), axial super-resolution imaging of organelle dynamics in a living biological sample, cortical astrocytes, an important type of brain cell. In the later case, our approach removes uncertainties in the interpretation of the nanometric axial dynamics of fluorescently labeled vesicles. Our technique is cheap, versatile and it has obvious applications in microscopies, profilometry and optical nano-metrology.

## INTRODUCTION

Ultra-thin polymer films play crucial roles as transparent spacer materials for the characterization of evanescent fields^1,2^, as transparent microwells^3^, or as scaffolds in microfluidics^4,5^ Thin films are also essential building blocks for optical coatings, organic/inorganic^4^ or photonic devices^5,6^, nano composites^7,8^, and for transparent nano- and micro-structures^3,9^. The fabrication of such ultra-thin polymer films goes along with their detailed characterization in terms of refractive index (RI), thickness Δ, (and, related, smoothness), as well as their uniformity and optical homogeneity. These different parameters are typically being quantified through a multitude of measurements involving a range of techniques, which include ellipsometry, stylus profilometry, atomic force microscopy (AFM), scanning electron microscopy, to verify the quality of the coatings.

In the current work, we show that - instead of characterizing thin transparent layers through a combination of complicated, expensive and time-consuming techniques - much of the same information can be retrieved through an all-optical readout, in a single measurement, on a standard inverted microscope equipped with a high numerical-aperture (NA) objective. This can be achieved by depositing a thin fluorophore layer on top of the transparent nanometric film to be characterized, and by measuring the fluorophore radiation pattern by aid of a phase telescope (Bertrand lens). The later shifts the focus from the front focal plane, i.e., the usual sample plane (SP), to the objective’s back focal plane (BFP), located inside the objective lens. BFP imaging allows us to collect the angular distribution (*k*-space representation, Fourier image). Quantitative BFP-image analysis thus reveals information that is otherwise lost^10,11^ and it allows us monitoring subtle changes in the fluorophore radiation pattern in real time.

Our technique is based on the fact that fluorophores located in a non-homogenous medium close to an interface display a directional emission different from the classical dipole emission in free space. Moreover, if these molecular fluorophores are located close enough for their near field to interact with the interface, they emit into angles otherwise forbidden by Snell’s law - a phenomenon known as supercritical angle fluorescence (SAF) or leakage radiation^12,13^. We exploit that this radiation pattern is further modified by changes in the proximity of the fluorophores to the interface, but also by the RI and homogeneity of the transparent film. We^14^ and others^15^ have shown that the precise position of the intensity maximum at the emission critical angle *ϑ*c provides a sensitive measurement of the RI of the medium in contact with the flurophores, while the relative intensity emitted into supercritical- and under critical angles encodes the fluorophore distance from the interface^16–19^. By collecting the information from the extreme periphery of the objective’s back pupil, in the region corresponding to *ϑ*_NA_ > d*ϑ* > *ϑ*c (where *ϑ*c are the critical angle given by Snell’s law and *ϑ*NA the limiting polar angle set by the NA, respectively), we can retrieve in the far-field the information originating from the evanescent wave emitted by the dipoles close to the interface.

In the present work, we first report of fabrication of a set of test samples in which a 5-nm homogenous layer of fluorophores (H6TPPS) is embedded in a thin polymer transparent film with RI of 1.33, at controlled axial distances from the surface, over a range of 10 to 340 nm. By imaging the BFP rather than the sample plane, we retrieve information from this near-field region. We measure the axial distance of the fluorophores at the interface, the RI of the transparent thin film and detect its local defects, when present. Our transparent medi^-^ um has a refractive index of 1.33 (My-133-MC polymer) to mimic a cell environment for biological applications. Working with this polymer, and fabricating thin homogenous films is a challenge by itself and it is important for biological studies.

Our fabricated samples and SAF-based thin-film characterization have a large range of potential applications in nanotechnology, material science, molecular plasmonics, physics and nanobiology. In the current work, we describe its use for axial optical nano-metrology, and we report on an application for biological axially resolved nanoscopy. The here presented approach is easy to implement and it presents a significant advance over existing techniques, in terms of speed, cost and ease. We expect it to become a routine step in thin-film and 2-D material characterization, and a meaningful complement to evanescent-wave and point-spread function engineering based axial optical techniques, where it can considerably enlarge the information content compared to that of fluorescence-intensity images alone.

## RESULTS AND DISCUSSION

### Fabrication of samples with controlled ultra-thin polymer films of RI = 1.33

My-133-MC is a commercial, transparent polymer having a RI of 1.33, similar to that of the aqueous environment of the cytosol of a biological cell^20^. My-133-MC cures upon exposure to ambient moisture. Using this polymer, we assembled transparent layered structures through a combination of spin-coating and plasma surface-activation techniques. **Fig. 1a** shows a typical cross-sectional image, illustrating the characteristic nano-layered architecture of the resulting sandwiches. They exhibit, from bottom to top, a borosilicate substrate (BK-7 optical grade coverslip), a flat My-133-MC spacer layer of controlled nm-thickness; a 5-nm thin fluorescent-emitter layer and, on top, another transparent index-matched polymer-capping layer. This capping layer ensures that the dye molecules are exposed to a homogenous RI microenvironment. In the particular sandwich shown, we deposited additional conductive layers at the very bottom and on top, only for the purpose of electron microscopy. On the focused-ion beam (FIB) cross-section (*left*), we observe a remarkable homogeneity, smoothness and flatness of the different component layers (see also **Fig. S1** in the **Supplementary Material Online**).

**FIG. 1.**
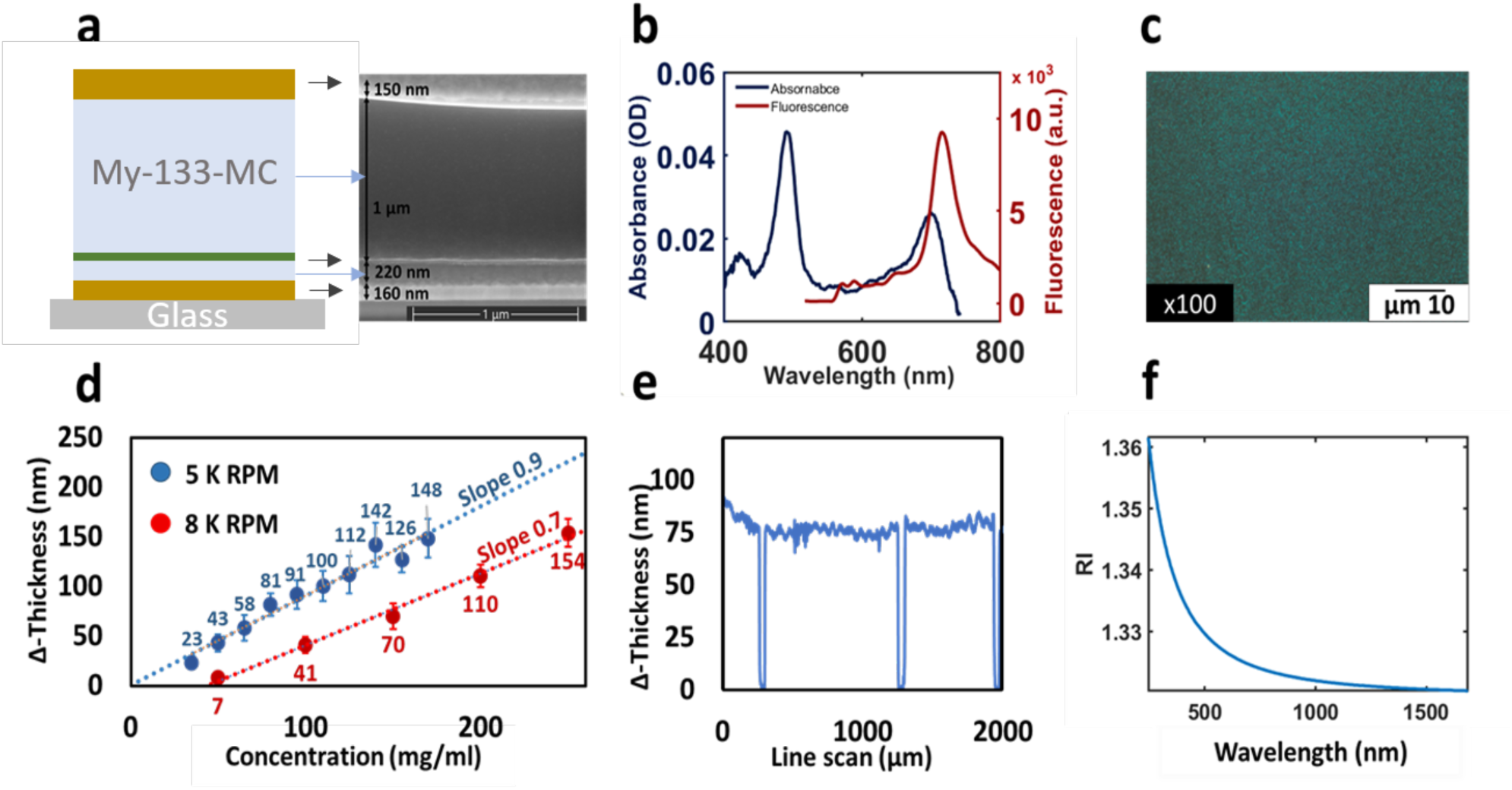
Multi-layered sandwich of transparent and fluorescent nanometric films with controlled properties. (a) schematic representation (*left*) and FIB cross-section (*right*) of a fabricated sandwich, alternating metallic (gold), My-133-MC polymer (light blue) and dye (green) layers. Metal layers were only added for FIB imaging. Note the thin fluorescent (4-nm) layer of H_6_-TPPS J-aggregate, and the well-defined layering. (b), absorbance (*blue*) and fluorescence (*red*) spectra of TPPS films sandwiched between polymer spacer and capping layers. (c), fluorescence micrograph of the dye layer showing a uniform concentration over large scale. (d), varying spin-coating parameters, ultra-thin layers of thicknesses down to a few nm thickness could be produced. Thickness was measured by Stylus profilometry, (e), that also provided a RMS roughness estimate of 1-2 nm. (f), Ellipsometry of the polymer layers, showing a RI around 1.33 in the visible range.

As an emitter we used a far-red emitting porphyrin derivative, (H6TPPS) J-aggregate, studied thoroughly by us before^21,22^. TPPS aggregates retained their bulk fluorescence properties when sandwiched between the two polymer layers, with neither a change in peak absorption (S2 at 491 nm, and S1 at 700 nm) nor in peak fluorescence (720 nm), **Fig. 1b**. From the measured optical density, we can estimate an equivalent dye concentration of “0.1M within the thin layer. Using our previously developed nitrogen-flow technique^21^ a large uniform dye coverage onto surfaces up to several mm^2^ can be obtained, **Fig. 1c**. We deposited these fluorescent films of about 5 nm thickness onto different surfaces, either the bare substrate, or an intermediate polymer layer, resulting in a well-defined nm fluorophore height Δ above the substrate. Spin coating allowed us to form a homogenous spacer of MY-133-MC and to systematically control its thickness, **Fig. 1d**. Different angular frequency and polymer concentration resulted in transparent layers of variable and precisely controllable thickness, ranging from a few to hundreds of nanometers. We used stylus profilometry and AFM for measuring the layer thickness, Δ, **Fig. 1e**. The resulting spacer layers had a RMS roughness (*R*q) of the order of 1-2 nm, independent of Δ. Ellipsometry confirmed a RI of about 1.33, as specified for My-133-MC polymer, **Fig. 1f**.

Taken together, through a combination of nano-fabrication and -characterization techniques we assembled and experimentally validated smooth, flat and uniform nano-layered structures alternating transparent and fluorescent layers. With their transparency and RI close to that of a biological cell, they feature a thin, homogenous and uniform dye layer, at a precisely controlled distance from the glass substate’s surface and can serve as a calibration tool for various axial super-resolution microscopies^2,23^.

### Thin-layer refractometry using far-field light

The radiation pattern of an emitter changes when a fluorophore approaches an interface. Near the boundary, the evanescent near-field emission component of a molecular dipole can couple to the interface and become propagative. By analogy to the excitation evanescent field in TIRF that requires supercritical illumination angles, this near-surface emission must propagate at supercritical angles, forbidden according to Snell’s law for far-field dipole emission. As a consequence, near-interface fluorophores emit an important power fraction into the higher-index medium (*n*2), predominantly beyond the emission critical angle *ϑ*c=asin(*n*1/ *n*2). Analyses of supercritical angle fluorescence on pupil plane images can retrieve *ϑ*c and from that the RI of the local environment of the fluorophores, *n*1 = *f*/*rc*, where *rc* is the equivalent critical radius and *f* the focal length of the objective (see Fig. S2 and S3 in **Supporting experimental procedures, online)**^14,15^

We implemented BFP imaging on a custom combined total internal reflection and supercritical angle fluorescence (TIR-SAF) microscope built from optical bench components (**Fig. S2**). Our instrument allows for the continuous selection of the polar illumination angle at the sample, between *ϑ*=0° (epifluorescence, EPI) and about ±74° (TIRF). We limited the scan angle to this value so as not to clip the laser beam by the measured effective NA*eff* of 1.465 (∼ 75.7°). Before each experiment, we systematically ran a test protocol validating alignment, setup performance and the absence of coverslip tilt (**Fig. S3**). In the following experiments, the emission from a 5-nm thin J-agg layer was bright enough for recording highly contrasted BFP images on a standard sCMOS camera with reasonable acquisition settings (*P* = 40 µW at the objective and 100-ms integration time), **Fig. 2a**. The recorded BFP images revealed a radially symmetric emission, indicative of non-preferentially oriented dipoles. They also showed the expected characteristic intensity transition at the emission critical angle *ϑ*c with an intensity maximum just above *ϑ*c, and an outer boundary corresponding to Na*eff* of the used objective^14,24^. Line profile intensities, but not peak positions and measured RI values, were sensitive to coverslip tilt (**Fig. S4**). The deposit of My-133-MC polymer on top of a dye-coated BK-7 substate (*Δ* = 0 nm), changed the measured BFP intensity profile and RI from 1.047 ± 0.002 (air) to 1.3182 ± 0.003 (My-133-MC), **Fig. 2b** (see panel **c** for a polar-plot representation as radiation patterns). The BFP-image cross sections and the measured RIs were independent of the azimuthal orientation of the very line profile chosen (panel b). As a consequence, our SAF-based RI measurements were robust and highly consistent for independent samples, **Fig. 2d**.

**FIG. 2.**
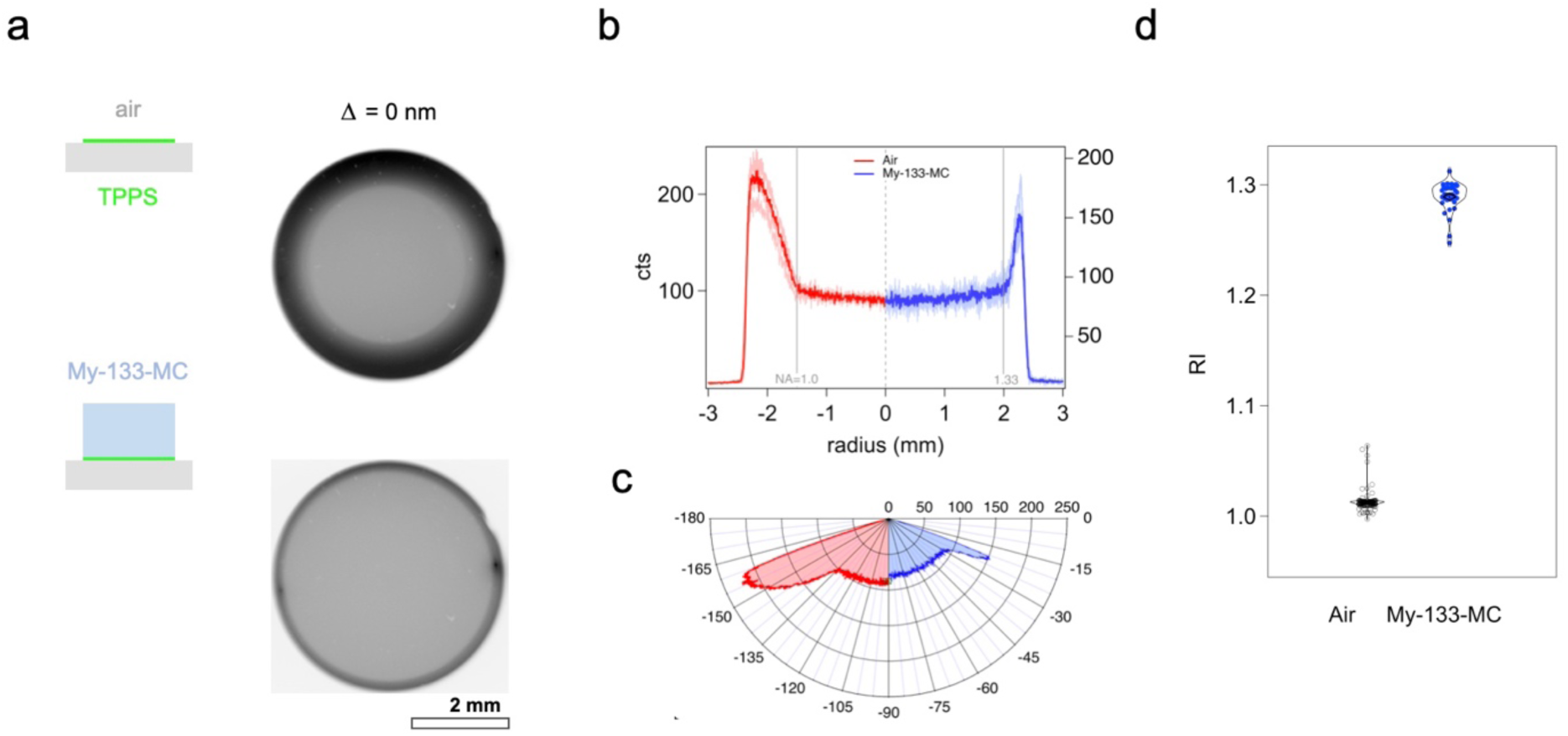
Supercritical-angle fluorescence (SAF) -based measurement of the refractive-index of ultra-thin transparent layers. (a) Schematic illustration of the sample (*inset*) and observed radiation patterns by BFP imaging of a 5-nm TPPS J-aggregate layer homogeneously deposited directly on the BK-7 substrate and left in air (*top*), or else covered a 3-µm thick My-133-MC polymer capping layer (*bottom*). Fluorescence excitation by TIRF. Image contrast inverted for better clarity (b), radial cross-sections of the images in (a), were largely independent of azimuthal *φ*. Thin, pale traces are pro-files measured at *φ* = 0°, 45°, 90° and 135°, respectively. Solid lines are azimuthal averages and are shown as polar plots in (c), revealing the characteristic directional lobes at supercritical angles. Note the change in emission pattern for the polymer-covered dye layer (blue). From the emission critical angle (grey in (b)), we obtained the refractive indices (RIs) of air and My-133-MC polymer, (d). Violin plots show population distributions over *n* = 129 (air) and 34 (polymer) measurements, respectively.

In order to characterize the uniformity of the produced My-133-MC polymer layers over larger spatial scales and to independently confirm our SAF-based RI measurements, we studied the polymer films by nanoplasmonics. Surface plasmons are evanescent waves that strongly depend on the dielectric function of the medium in contact. To avoid tedious phase-matching and to couple free propagating light to plasmonic modes, we turned to plasmonic structures that scatter the incoming light and overcome the *k*-vector mismatch by additional momentum from the reciprocal lattice. We milled periodic nano-hole arrays in a thin Au-film deposited on the BK-7 glass substrate. Under white light illumination we observed on a dark-field microscope structural colors that varied among plasmonic hole arrays according to their different periodicities, *P*, i.e., inter-hole spacing (340, 380 and 420 nm)^10,22,25^ **Fig. 3c** shows transmission spectra of those arrays upon white-light illumination. The same peaks appeared for plasmonic structures covered with either water or My-133-MC polymer, indicating the same RI for both media. Compared to the localized SAF-based RI measurement limited by the field-of-view of the x100 objective (Fig 2), this plasmonic approach intrinsically probes different locations of the polymer layer over >100 µm, indicating that the deposited My-133-MC layers have a homogenous RI over larger length-scales.

**FIG. 3.**
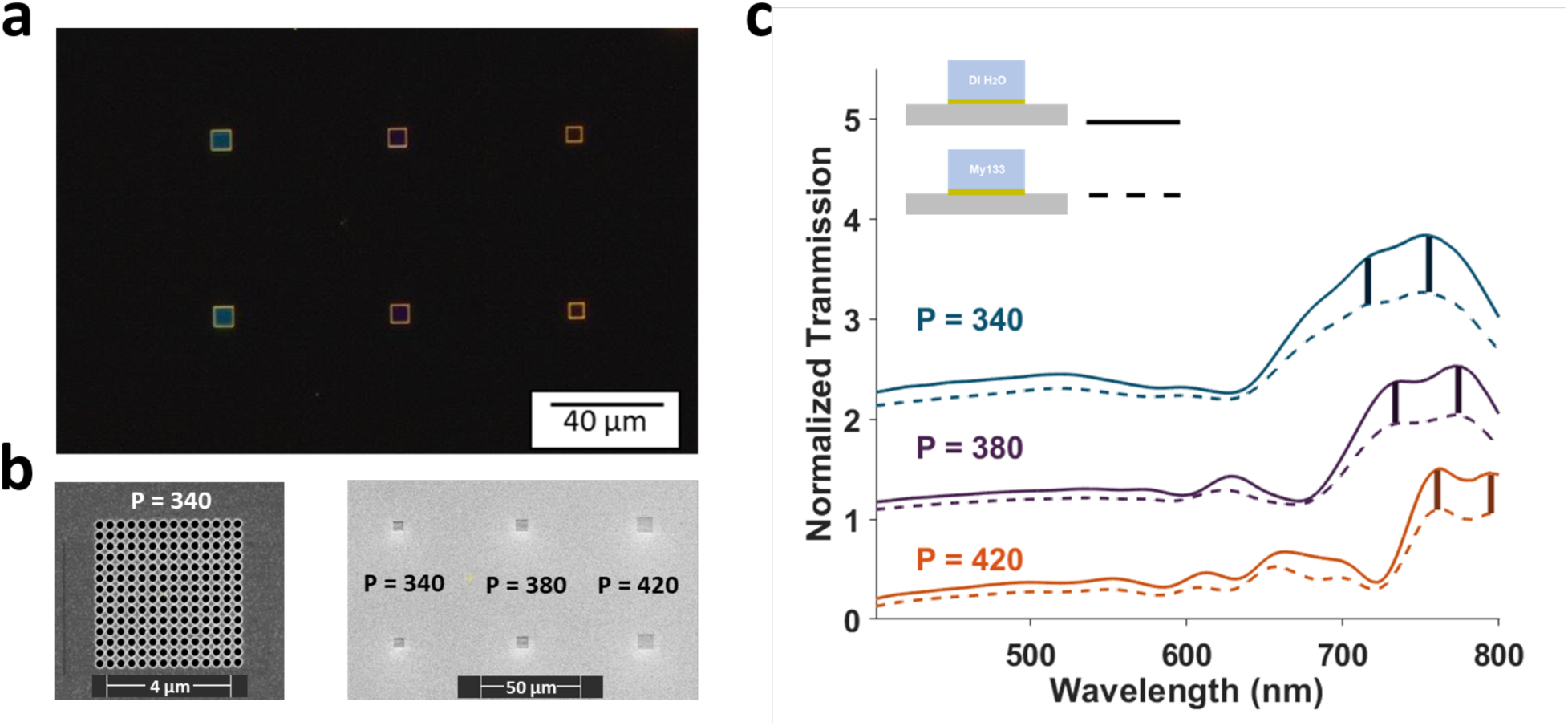
RI homogeneity in ultra-thin polymer layers. (a), dark-field image upon white-light illumination and, (b), electron micrographs of a single plasmonic hole array (*left*) as well as a larger plasmonic sample comprising several patterns with different periodicities *P* (in nm, *right*). (c), Normalized transmitted intensity vs. wavelength upon white-light illumination, showing similar responses of the plasmonic hole array when covered with water (*through* line) or My-133-MC polymer (dashed). Vertical lines highlight identical peak positions for the plasmonic modes either at interface with water or My-133-MC, respectively.

Taken together, our ellipsometric (fig.1f), SAF-derived (fig.2) and plasmonic RI measurements (fig.3) confirm a RI of ultra-thin My-133-MC layers close to that of water, validating this transparent polymer as a suitable optical medium for diverse biomimetic applications in fluorescence microscopy and in biophotonics.

### Detecting imperfections of the polymer thin layers

We filtered the polymer solution to remove air bubbles and obtain flawless, uniform and homogenous My-133-MC polymer films. Nevertheless, BFP images sometimes revealed a conspicuous double-ring structure featuring two intensity transitions, one at RI = 1.33 and another one at RI = 1, **Fig. 4a** (*left* half image). The inner ring was dimmer compared to the outer one, see *red arrow* on panel (a), and the diameter of the outer ring corresponded to that observed with the majority of produced polymer samples (*right*). We reasoned that such a double-ring pattern could result if some - but not all - fluorophores were exposed to air rather than the polymer. Such a non-homogeneous dye environment could result from cavities, cracks or air-bubbles in the polymer (**Fig. S6**). Indeed, a *post-hoc* FIB cross-section on these samples confirmed the presence of µm-sized bubbles in the polymer layer, **Fig. 4b**. The observed bubble size is probably an overestimate given that SEM images were acquired in vacuum. Imperfections could equally be observed by the light transmitted through the plasmonic hole arrays on which the polymer layers were deposited (**Fig. S7**), lending further plausibility to our interpretation. Dual RIs can be also appear when cracks or dimples were present. Fig. 4c shows such an example for which a marked RI change was observed upon wetting the polymer surface. Starting from an air-dominated environment, pipetting a drop of polymer onto the surface changed the RI – from 1 (*green*), to 1.3 (*blue*) - the change occurring at the moment of the deposit of the drop (*red*).

**FIG. 4.**
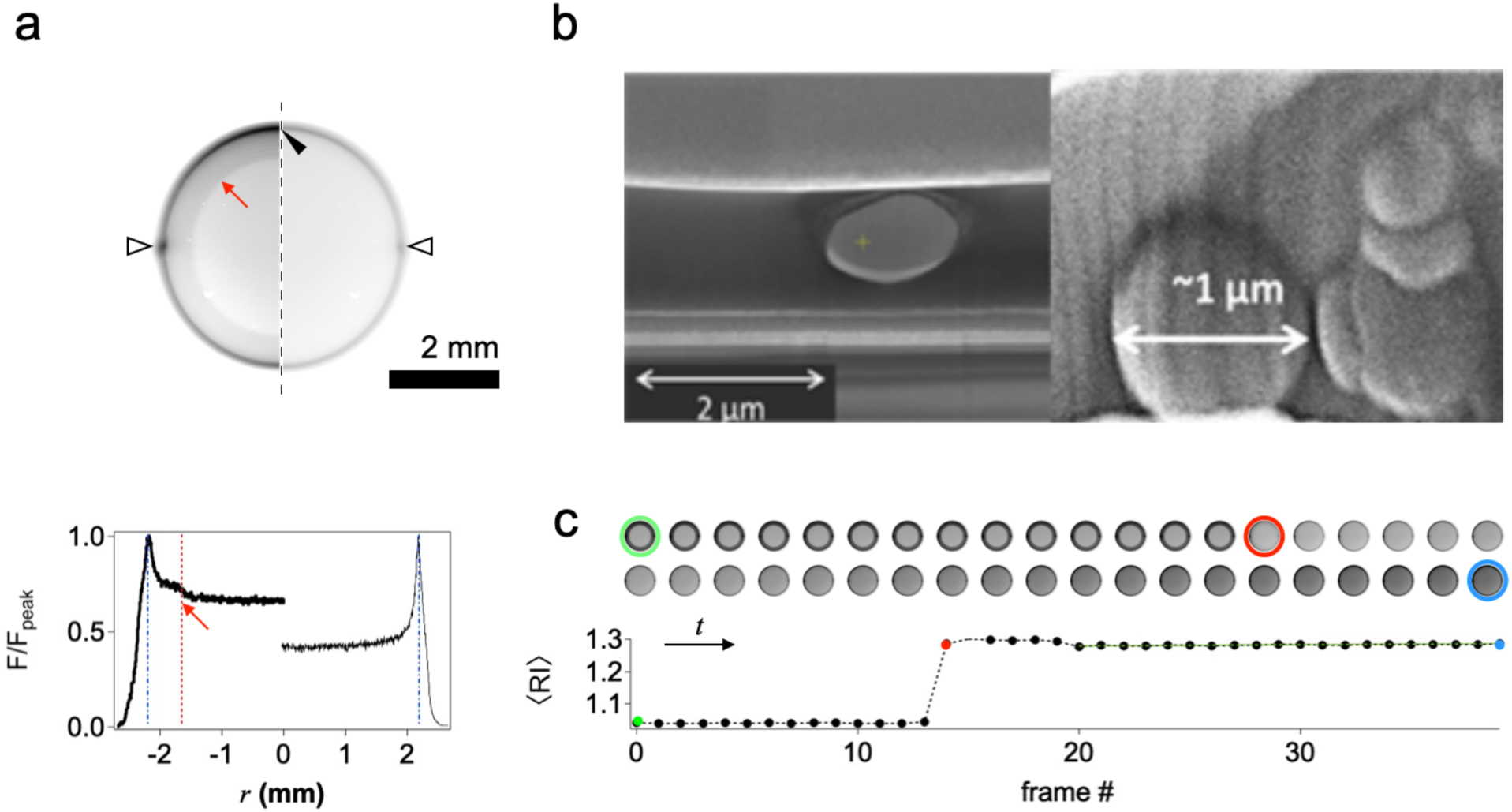
SAF-based detection of flaws in nano-fabrication. (a), *left* half, example BFP image of an imperfect polymer layer displaying a double-ring structure at RIs corresponding to both air (red arrow) and My-133-MC (black arrowhead). *Right*, single-ring structure observed for a flawless sample for comparison. Intensity of the left half image increased for better visibility, contrast is inverted, as in fig.2. *Bottom*, corresponding normalized-intensity line profiles. Note the little kink on the left profile (*red* arrow), absent on the right. Dash-dotted and dotted lines indicate RI 1.33 and 1, respectively. (b), two examples of *post-hoc* FIB cross-sections of imperfect polymer films. (c), example of a My-133-MC polymer layer with cracks. *Top*, time-lapse BFP-image series and SAF-derived RI, *bottom*. At frame #14, a drop of polymer was pipetted on top of the sample, increasing the measured RI from close to 1 to 1.33. Colored rings and spots, respectively, identify corresponding time frames.

Thus, the information contained in the fluorophore radiation pattern does not only permit precise thin-layer refractometry but it also reveals imperfections, the detection of which requires otherwise more involved, lengthy and expensive experiments. The time-series in **Fig.4c** furthermore illustrates that our technique permits time-resolved measurements and *in operando* refractometry, an important feature for sensors and thin-film devices. Based on this SAF-based BFP-image refractometry, we retained only samples that passed the quality gate of a single-ring BFP pattern, and we next investigated how changes in the thickness of the spacer layer, i.e., the fluorophore height above the substrate, modified the radiation pattern.

### Measuring axial nanometric distances from SAF/UAF ratios

Capitalizing on our ability to fabricate controlled thin nanolayers, we next studied sandwiches with different fluorophore-substrate distances Δ. We prepared otherwise identical sandwiches featuring the TPPS emitter layer at Δ = 0, 5, 15, 25, 32, 29, 75 and at 340 nm, respectively, three of which are schematized on **Fig. 5a**. We used TIRF, the focused laser spot is seen in the periphery of the BFP image for the incoming and back-reflected beam, to excite only near-interface fluorescence. As expected upon evanescent-wave fluorescence excitation, the intensity dropped with increasing Δ, **Fig. 5b**. While the evanescent-wave penetration depth can be calculated from the local RI of the medium (*n*_1_ = 1.33 for My-133-MC) and the incidence angle of the illuminating beam at the laser wavelength (that can be fitted from the laser spot observed on the BFP image^26^), the actual depth in a given experiment, on the particular microscope is much less certain. How deep the excitation light actually penetrates into the sample is difficult to know and is modified by sample, surface and objective imperfections^23^, which altogether make the quantitation of TIRF intensity images and the measurement of axial fluorophore height from single-angle TIRF data problematic.

**Figure 5.**
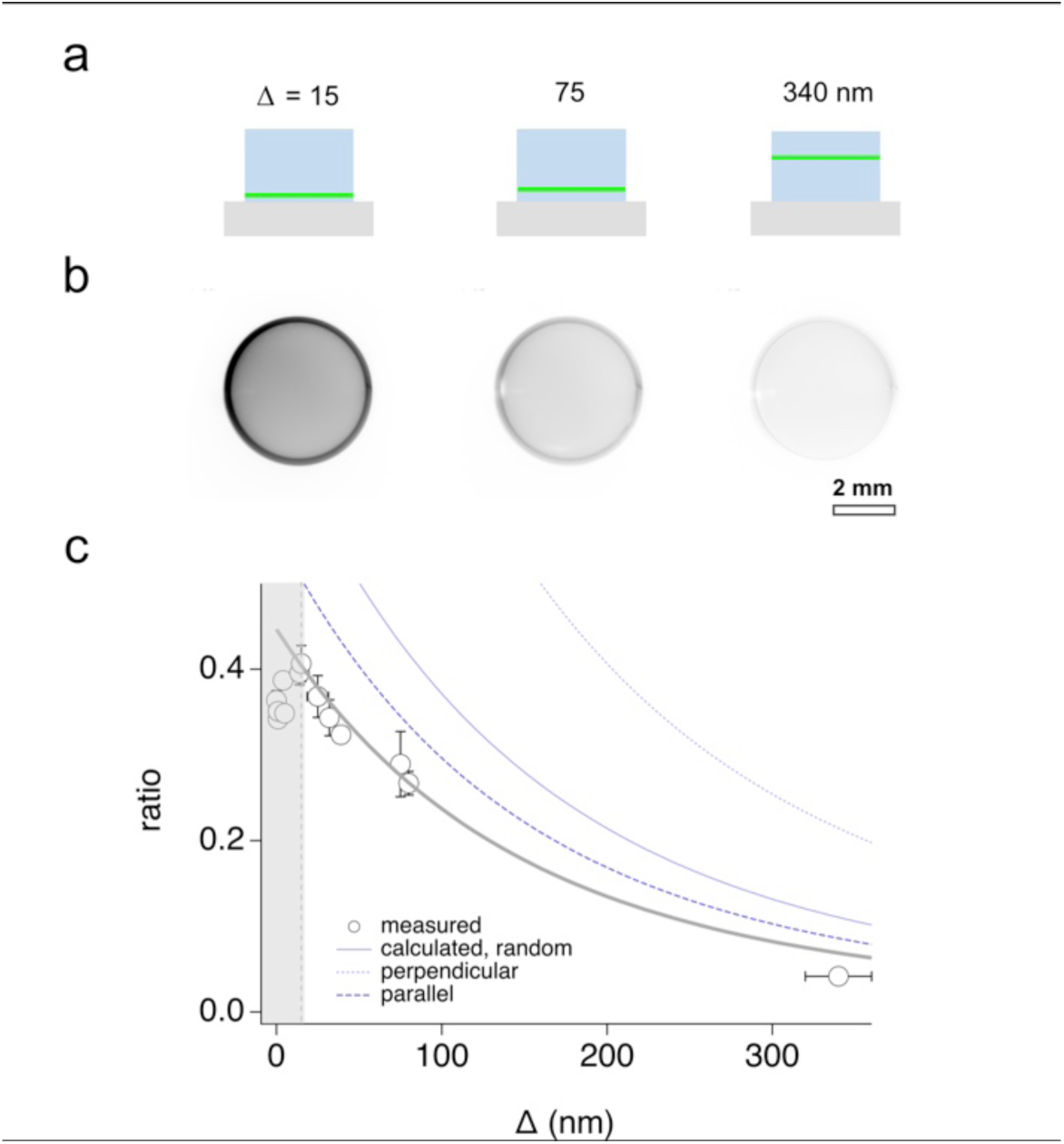
SAF-based axial profilometry. (a), schematic representation of a subset of three prepared test slides, featuring a thin TPPS J-aggregate emitter layer sandwiched at different fluorohpore heights Δ between a flat and homogenous My-133-MC spacer and capping layer. (b), corresponding example BFP images after background subtraction and scaled to equal min-max intensity for direct comparison. (c), plot of the SAF/UAF intensity ratio *R* of supercritical vs. undercritical-angle fluorescence emission components vs. fluorophore height Δ. Symbols are measurements from segmented BFP images as in (b), *thin* lines are theoretical values calculated using the constant-radiated power dipole of Hellen & Axelrod (1987) for dipole orientations as indicated by the line dash. Grey through line is the same curve as for a dipole oriented in the incidence plane, scaled to best fit and suggesting a lower collected fraction of SAF compared to UAF (seen main text). The shaded region of the first surface-proximal 15 nm was omitted from fit. Error bars were measured Rq roughness (for x) and generated (for y) by allowing for +/-0.05 RI-unit uncertainty, respectively.

We therefore turned to a ratiometric measurement by plotting supercritical (SAF) vs. undercritical fluorescence (UAF) emission components, *R* = *I*SAF/*I*UAF, as a function of fluorophore height Δ, **Fig. 5c**. Because the light passing through the supercritical zone of the BFP originates entirely from the fluorophore near field, the farther away from the interface the fluorophore resides, the dimmer will be the SAF component^17,27,28^. UAF on the other hand, is almost constant, with the exception of a small surface effect at distances very close to the interface. Advantageously, the SAF/UAF fluorescence ratio normalizes for fluorophore concentration, laser intensity and other factors that linearly influence the measured signal. We segmented background-subtracted BFP images into undercritical and supercritical regions, based on the previously measured NA*eff* and RI and the integrated intensities over these regions. **Fig. 5c** graphs the evolution with Δ of the SAF/UAF ratio *R* = *I*SAF/*I*UAF, along with simulations for different fluorophore orientations, using the constant total-radiated power model from Hellen & Axelrod^29^. We see a good agreement with theory, except for the values very close (Δ <15 nm) to the interface, where the observed ratios were systematically lower than the predicted one. While we can exclude systematic errors in the thickness of the spacer layers even at that scale from our earlier electron-microscopy controls, one possible explanation is that surface quenching affects SAF stronger than UAF, or that fluorophore orientation is modified in a surface-distance dependent manner, e.g., by surface charges. Alternatively the objective might not be as efficient in capturing high-angle (SAF) as capturing low-angle (UAF) light. In fact, theory assumes a flat collection efficiency over the entire solid angle covered by the NA, which might not be the case for the used objective lens (as already observed others, see, e.g.,^30^). Might that as it be, a systematically lower fractional SAF also explains why the best fit (*grey* trace) corresponds to theory (*blue*), up to a constant scaling factor. Another reason for this offset could be a generally higher UAF fraction than expected: for example, if My-133-MC polymer was even slightly autofluorescent, this would lead to a relatively higher UAF component (resulting from the spacer and capping layers) and be detectable under the low-light conditions of the TIRF experiment. Yet, fluorescence spectroscopy of bulk of polymer did not reveal any detectable autofluorescence. Of course, the objective itself or the immersion oil could fluoresce and contribute UAF, despite a detection above 720 nm. Taken together, even with the note of caution for very small values of Δ, SAF/UAF ratiometry allows a reliable, quantitative analysis of the thickness of nanometric layers in the sub-100-nm range, with a precision much higher than that of TIRF-intensity measurements alone.

### Biological axial nanoscopy using SAF/UAF emission ratios

One example of biological nanoscopic imaging that would benefit from a precise axial fluorophore localization is the analysis of single-vesicle mobility during regulated exocytosis. These experiments were pioneered in TIRF studies of the of the late 1990^31–35^ and have widely been used in the cell- and neurobiology community since. Exocytosis involves the transport, docking and release of tiny, membrane-delimited and cargo-loaden compartments (‘vesicles’) that are formed and filled inside the cell, transported to the cell’s outer membrane and that, upon a stimulus - often a rise in the intracellular free calcium concentration ([Ca2+]i), fuse with the outer membrane to release their contents, see **Fig. 6a**. In such experiments, it would be important to know if a vesicle is close to the membrane or just a bit above, if it moves towards it or away from it, or if it simply loses fluorescent content by leakage or bleaching - questions that cannot be easily answered from an intensity TIRF recoding, due to the unknown evanescent-wave penetration depth and the confounding effects of dye concentration and fluorophore distance on single-vesicle brightness. Unfortunately, the fast dynamics of the process of exocytosis does neither permit to capture *z*-stacks of images for axial localization at higher precision, nor to acquire a variable-angle series of TIRF images for a ‘tomographic’ reconstruction of vesicle distance.

**Figure 6.**
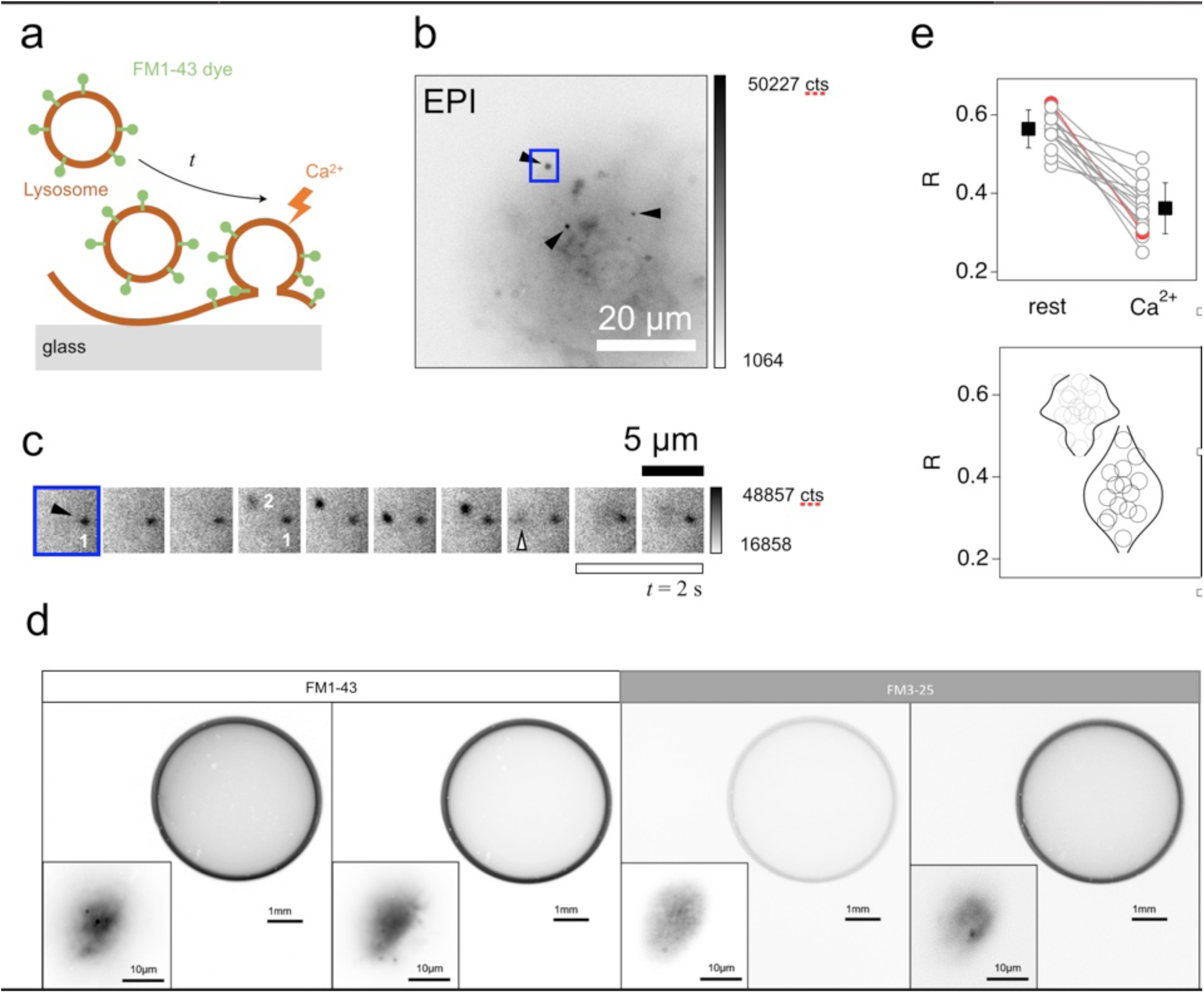
SAF-based disambiguation of TIRF image data. (a), schematic representation of the intracellular steps characteristic for calcium (Ca^2+^) -regulated exocytosis. FM dye (*green*) is a membrane-resident (brown) amphiphilic fluorophore that is internalized into cortical astrocytes and labels a tiny intracellular vesicular compartment called lysosome. These organelles approach the basal plasma membrane of cells cultured on a glass coverslip and are released in a Ca^2+^-dependent manner (arrow). (b) epifluorescence (EPI) image showing a diffuse cytosolic labeling as well as bright spots representing individual lysosomes (arrowheads). (c), zoom on the rectangular region shown in blue on (b). The time-lapse image series acquired at 1 Hz shows two lysosomes, “1” and “2”. While 2 (solid arrowhead) is present from the beginning to the end of the recording, 2 arrives in frame #3, approaches the membrane and releases its content (“exocytosis”) at the moment identified by an open arrowhead. The corresponding axial re-distribution of fluorophores is seen in (d), which examples of BFP images of ROIs (*inset* images) of basal membrane in cultured cortical astrocytes upon TIRF illumination, with either FM1-43 (*left*) or the spectrally identical but slightly more hydrophilic FM3-25 dye, *right*. Note the change in SAF intensity for ROIs showing membrane proximal lysosomes compared to void regions. The *rightmost* image is an exception because the SAF/UAF intensity ratio *R* dropped despite an organelle still visible, seen as a spot on the inset SP image. (e), Comparison of *R* values before (rest) and after (Ca^2+^) stimulation, showing a systematic trend towards lower ratios, indicative of a loss of near-interface fluorophores consistent with exocytosis The red trace follows the trend but corresponds to the rightmost image on d, where the lysosome apparently moved back into deeper cytoplasmic regions, thus lowering ISAF and hence *R*.

We therefore studied, in cultured mouse cortical astrocytes labelled with an amphiphilic styryl pyridinium dye^36^, the radiation pattern emerging from small regions of evanescent-wave illuminated membrane, close to the coverslip. In astrocytes, a glial type of brain cells that are neighbors to most neuronal synapses, the green-yellow emitting non-membrane permeable dyes FM1-43 or FM3-25^37^ show a behavior and uptake-mechanism different from that in neurons and are internalized to label larger vesicular compartments, called lysosomes^36,38^. On fluorescence images these lysosomes appear as diffraction-limited fluorescent spots in front of a diffuse background, **Fig. 6b**. On a timelapse series of TIRF images acquired close to the bottom of the cell, we can appreciate the movement of some lysosomes in and out of the evanescent field, while others dwell patiently close to the plasma membrane, **Fig. 6c**. Stimulation of the cell by application of extracellular adenosine triphosphate (ATP) activates purinergic receptors and increases the intracellular Ca^2+^ concentration, leading to a loss of most near-membrane lysosomes. This interpretation is confirmed by analysing BFP images that were taken alternately with sample-plane images and that reproducibly displayed *R* values of 0.59-0.63 (0.56 ± 0.05, *n* = 15) for cells with neither exocytic events nor overt mobility, **Fig. 6d**, *leftmost* and *left* panels. Following the loss of near-membrane lysosomes, the measured SAF/ UAF ratios were consistently lower, *R* = 0.36 ± 0.06, explained by an average localization of the remaining dye molecules deeper within the cell, more distant from the interface. Clearly, the remaining FM dye molecules less capable of emitting SAF, **Fig. 6d**, second panel from *right*. The interest of concurrent sample-plane and quantitative BFP imaging is evident from the rightmost panel where *R* was 0.30 ± 0.02, despite the fact that a single lysosome was still present on the sample-plane image. While pure TIRF intensometry would suggest a membrane-proximal location, SAF/ UAF ratiometry revealed a *R* value of around 0.3 and hence an apparently greater distance from the basal cell membrane than suggest from the intensity reading alone. SAF/UAF ratiometry can thus eliminate unknowns like dye concentration, vesicle diameter or local illumination heterogeneities^39^ and permit a more reliable interpretation of near-membrane fluorescence.

By comparison with the ratios of fig.5c, we can infer that the lysosomal dynamics occurs in near-membrane region at the bottom of the cell, in a region closer than 100 nm to the coverglass. SAF/ UAF ratiometry thus allows the resolution of axial fluorophore dynamics within this range. One limitation of the *R* measurements in the current implementation of the microscope, however, is that they average fluororophore properties over the entire field-of-view (some 40 µm diagonal). More resolved measurements can be expected when reducing the size of the illuminated field-of-view, an approach that we currently implement using Bessel-beam illumination with an evanescent needle of light^40^.

Very clearly, our proof-of-concept biological experiments illustrate the usefulness of concurrent sample- and back-focal plane imaging for interpreting and disambiguiting the dynamics of biological fluorescence, and they also illustrate the interest of fluorescence imaging with combined TIRF excitation and SAF detection^13,41^ in conjunction with a rapid, online BFP-image analysis. We expect our technique to have applications in various fields of “surface biology”, like the dynamics of ER-PM junctions that link the endoplasmic reticulum to the plasma membrane, cellular adhesion sites, or biofilm growth, but also in surface chemistry and material science.

## Conclusion

Taken together, the combination of nano-fabrication and -characterization techniques allowed us to assemble and validate controlled, smooth and uniform nanometric thin fluorescent transparent layers of precisely defined thickness and having a RI close to that of a biological cell. Our sandwich samples can serve, e.g., as a nanoscopic axial ruler for axial super-resolution microscopies and they provide a valuable reference for interpreting the axial dynamics of intracellular processes in the near-membrane space of live cells, cultured on a glass coverslip. We demonstrate that combined sample-plane and quantitative BFP imaging provides unique insight into fluorophore properties and dynamics. Particularly, we show that SAF based refractometry and SAF/ UAF ratioing can be used for exploring the physical and chemical microenvironment of fluorophores and can be used for time-lapse *in situ* imaging of fluorophore dynamics at or close to a dielectric interface.

BFP imaging is easily implemented on a standard inverted microscope and it allowed us to retrieve axial fluorophore information in the sub 100-nm range, SAF/UAF ratios provide complementary information to fluorescence intensity. In fact, the analysis of the fluorophore radiation pattern gives access to the RI of the fluorophore-embedding medium, the fluorophore height (or film thickness) and orientation of emitters, with the only condition that they are sufficiently close for their near-field and SAF emission component to be modulated by the surface proximity. As a simple and reliable far-field optical technique, the SAF-based thin-film characterization is an interesting alternative to more complex, time-consuming and expensive techniques. We see additional potential in the field of molecular plasmonics, or polaritons in chemistry, where the nanofabricated samples and the proposed k-space imaging technique togehter offer a powerful tool for studying strong coupling between emitters and plasmonic system ^22,25,42,43^. In fact, for the here reported controlled nanolayers, the interaction between molecules and plasmons is very well defined, and so is the coupling (Rabi splitting) that is not averaged over emitters at different distances. Finally, we also expect axial-nanoscopic rulers to become a standard for various fields areas of nano-metrology. With their transparency and RI close to that of a biological cell, they feature a thin, homogenous and uniform dye layer, at a precisely controlled distance from the glass substate$s surface and can be used as calibration standards for axial optical sectioning, super-resolution and fluorohpore localisation techniques. Finally, the robust SAF/UAF ratios can instruct axial measurements from unknown samples, e.g., to disambiguate biological recordings from TIRF microscopes that have been challenging to interpret.

## EXPERIMENTAL PROCEDURES

### Materials

We chose My-133-MC (MyPolymers LTD, Ness Ziona, Israel) for its RI close to that of water and two fluorinated (Hydro-Fluoro-Ether, HFE) compounds as solvents for this polymer, Novec™ 7500 or 7100 (3M^TM^, Minnesota, USA). For the fluorescent emitter layer, we opted for H6TPPS4 (meso-tetra 4-sulfonatophenyl porphyrin) (Sigma-Aldrich, #88074) in its J-aggregate form due to its far-red/near-IR emission remote from autofluorescence and most biological fluorophores (peak absorption/emission wavelength, *λ*abs/ *λ*em = 488/720 nm^21^).

### Thin-film polymer deposition

*#*1 BK-7 glass coverslips (Menzel Gläser, Braunschweig, Germany) were cleaned in 1/100 Hellmanex/double-distilled-deionized water (DI water, 18.2 mΩ) and sonicated (20 min, 40 °C). They were then rinsed thoroughly with DI water, ethanol, and DI water again (30 s each). Substrates were dried with a N_2_ flow and immediately used thereafter. MY-133-MC polymer solution was filtered and thin films were spin-coated (Laurell, WS-650) and cured under ambient conditions (RT, 20-22°C) during at least 6 hours. My-133-MC polymer resin was applied either pure (undiluted) or diluted in HFE-7500.

### Thin dye-layer deposition

H_6_TPPS_4_ was dissolved in DI water and aqueous nitric acid (65% w/w, Sigma-Aldrich) to reach pH∼ 1. The thin film-deposition procedure is described elsewhere^21^. Absorption and fluorescence spectra were recorded both from films and in solution, using a home-built microspectrofluorimeter built around an Olympus inverted microscope coupled to a spectrophotometer (IsoPlane SCT-320, Princeton Instruments) and equipped with a charge-coupled device camera (CCD, PIXS1024b, Princeton Instruments).

### Nanoplasmonics

Plasmonic structures were milled in our in-house clean rooms, using a Gallium focused-ion beam (FIB, Helios NanoLabDualBeam 600, FEI) into previously deposited 200-nm thin silver films. The fabrication process was entirely calibrated for milling depth, beam size, and beam deflection. The integrated scanning electron microscope (SEM) enabled the *in operando* imaging of the fabricated structures as shown.

### Thin-film characterization

Film thickness and roughness were measured using a Stylus Profiler (DektakXT, Bruker) in at least 6 different areas. RMS surface roughness (*R*_q_) was calculated using Mountains8, Digital Surf. Results were cross-validated on the same specimens with AFM (Nanoscope, Veeco). The AFM images are shown as pseudo-colored height maps, from which *R*_q_ was calculated. RIs were measured by spectroscopic ellipsometry (J.A. Woollam, M-2000), nanoplasmonics and SAF (see below). For ellipsometry, we constructed a model for the My-133-MC polymer, based on the known wavelength and thickness of the deposited layers, obtained by SEM (See **Fig. S1**).

### Combined TIR-SAF imaging

Optical microscopy was performed on a custom set-up assembled from optical bench components (see **Fig. S2** as well as **Supporting Experimental Procedures** for details). Briefly, the beam of a 488-nm laser (Coherent Sapphire SF 488-50) was cleaned up with a 488-nm notch filter, attenuated with neutral density filters, spatially filtered and expanded to “2” diameter. It was then scanned by a rotary mirror and tightly focused in the BFP of a ×100/1.46NA objective (a Plan-Apochromat, oil DIC M27, ZEISS). Images were acquired either upon evanescent-wave excitation (with the spot positioned in the extreme periphery of the objective’s BFP) or upon EPI excitation (spot on axis, at 0, 0) using the laser powers and exposure times as indicated. Fluorescence was collected through the same objective, filtered by a ultra flat (2-mm) long-pass 488LPXR dichroic mirror (AHF Analysentechnik), RET493LP (Semrock) and ET520LP (Chroma) long-pass filters. Band-pass filters were housed in independent filter wheels and used as stated in the figure legends). The filtered fluorescence was imaged on a back-illuminated sCMOS camera (PCO edge4.2bi). The effective pixel size in the sample plane was (30 ± 1 nm/px, allowing for a 3-to 4-fold binning. A Bertrand lens mounted on a motorized flipper (Thorlabs) allowed shifting the focus to the objective’s BFP (3.36 µm/px on BFP images).

### SAF/UAF ratio image analysis

SP and BFP fluorescence images were subtracted with their respective dark images. BFP images were segmented into SAF and UAF regions in a multi-step process: we first found the center of the BFP pattern and the number of pixels corresponding to the limiting radius of the NA (*r*_NA_). For this, we binarized the image and searched the white disk on the black background by an object recognition algorithm implemented in MATLAB. Next, we calculated the pixel radii corresponding to the critical angle of the pattern (*r*_c_) by imposing an *a priori* RI estimate from independent measurements, *r*_c_ = *r*_NA_.RI/NA. Finally, we generated error bars by allowing for a positive and negative deviation (dRI) according to *r*c^(±)^ = *r*_NA_.(RI±dRI)/NA. Together, the center coordinates, *r*_NA_ and *r*_c_ enable us to segment the image into three areas: background, SAF and UAF. With the background subtracted, we integrated the intensity over each area and calculate the SAF/UAF ratio *R* = *I*_SAF_/*I*_UAF_, where *I* is the respective integral over SAF and UAF areas. Repeating this integration for each dRI gave us an estimate of the dependence of SAF/UAF ratio on the accuracy of the RI estimate. For the error bars shown we opted for dRI = ±0.05.

### Cell culture, staining, and microscopy

Cortical astrocytes were prepared from newborn mice, cultured and labeled previously described^36^ (See **Supporting Experimental Procedures** for details). We systematically applied 100 µM ATP at the end of the recordings to test for cell viability and raise the intracellular Ca^2+^. Labelling and the overall aspect of the cell culture was confirmed over a larger field on a custom upright microscope fitted for EPI and prism-type TIRF microscopy^44^. Fluorescence was excited at 488- and 568-nm wavelength from a polychromatic tuneable light-source (TILL Photonics). After appropriate filtering, fluorescence was detected on a EMCCD camera (Photometrics). METAMORPH and ImageJ software were used for acquisition and image analysis.

For combined TIR-SAF imaging, the same cells were transferred to the home-built inverted TIR-SAF microscope ^41,44–46^ and a tiny sub-region of the cell’s footprint containing FM-loaded lysosomes imaged. Equipment and imaging parameters were similar to that described above.

## Acknowledgements

We thank Daniel Azulai for help with the polymer deposition and Naftali Kanovsky and Prof. Shlomo Margel for advice regarding the polymer. Thierry Bastien provided custom fine mechanics. This study was financed by the CNRS, University Paris Cité, and - mostly - by the European Union (H2020 Eureka! Eurostars, ‘NanoScale’, E!12848, https://nanoscale.sppin.fr, to MO and AS). AS acknowledges support from the French Ministry of Foreign Affairs and the French Embassy in Tel Aviv (Chateaubriand fellowship). The authors are grateful for mobility support from a Franco-Israeli CNRS-LIA, ‘ImagiNano’, and the FranceBio-Imaging large-scale national infrastructure initiative (FBI, ANR-10-INSB-04, Investments for the future). The Oheim lab is a member of the C’Nano Excellence Network in Nanobiophotonics (CNRS GDR2972).

