## Supplementary Material for "Characterizing nanometric thin films with far-field light"

This SI contains

A list of abbreviations page *i*

6 supporting figures page *ii - vii*

Supplementary experimental procedures page *ix - x*

### LIST OF USED ABBREVIATIONS

| | | |
|---|---|---|
| AFM | - | atomic-force microscopy |
| ATP | - | adenosine triphosphate |
| BFP | - | back-focal plane |
| BK-7 | - | borosilicate crown optical glass |
| DIW | - | de-ionized water |
| EMCCD | - | electron-multiplying charge-coupled device |
| EPI | - | epifluorescence |
| EW | - | evanescent wave |
| FIB | - | focused ion beam |
| FM1-43 | - | Pyridinium, 4-[2-[4-(dibutylamino)phenyl]ethenyl]-1-[3-(triethylammonio)propyl]-, dibromide (CAS: 149838-22-2) |
| My-133-MC | - | moisture-cured coating material, with a refractive index of 1.33 |
| NA | - | numerical aperture |
| ND | - | neutral density |
| RMS | - | root-mean square |
| RI | - | refractive index |
| RT | - | room temperature (20-23°C) |
| SAF | - | super-critical angle fluorescence |
| sCMOS | - | scientific complementary metal oxyde sensor |
| SIM | - | structured-illumination microscopy |
| SP | - | sample plane |
| STED | - | stimulated emission depletion |
| TIRF | - | total internal reflection fluorescence |
| TPPS | - | sulfonatophenyl-tetraphenyl porphyrin |
| UAF | - | under-critical angle fluorescence |

#### SUPPLEMENTARY FIGURES

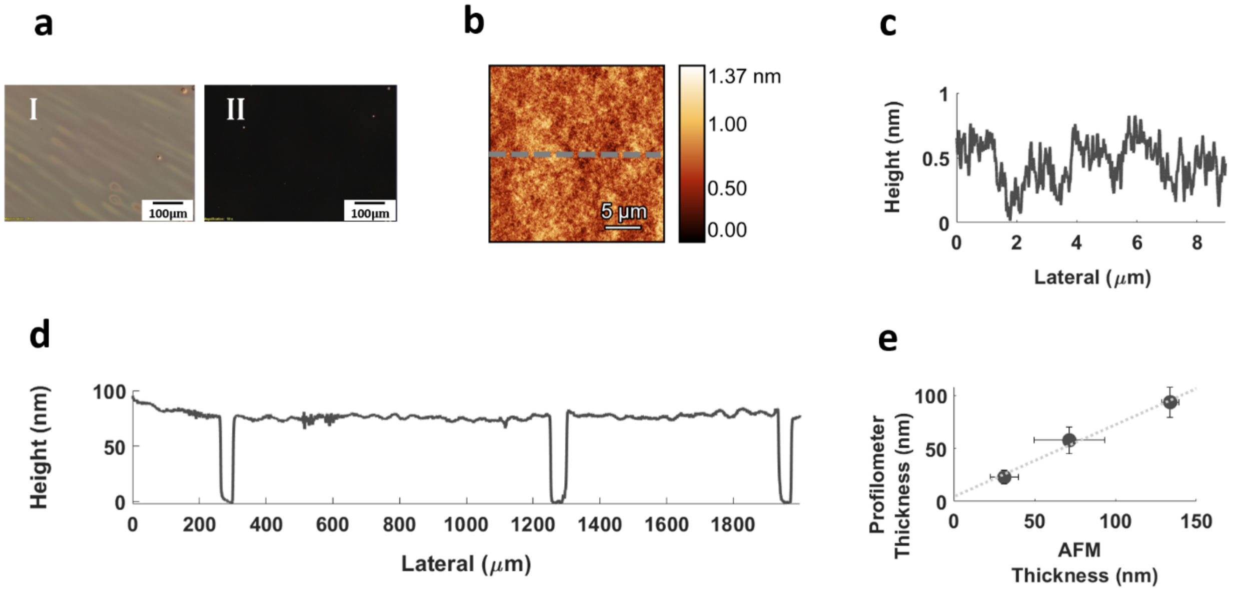


**Fig. S1.** *Additional characterization of My-133-MC-TPPS sandwiches* **(supplement to Fig.1)**. (**a**), phase (I) and dark-field images (II) show the absence of large-scale inhomogeneities and scattering centers, respectively. Scale bar is 100 µm. (**b**), example AFM micrograph and height profile, (**c**), of the resulting layer. (**d**), Example of stylus profilometry showing the uniformity and flatness over large areas. Kinks a scratches for height measurements. (**e**), Comparison of stylus profilometry and AFM results.

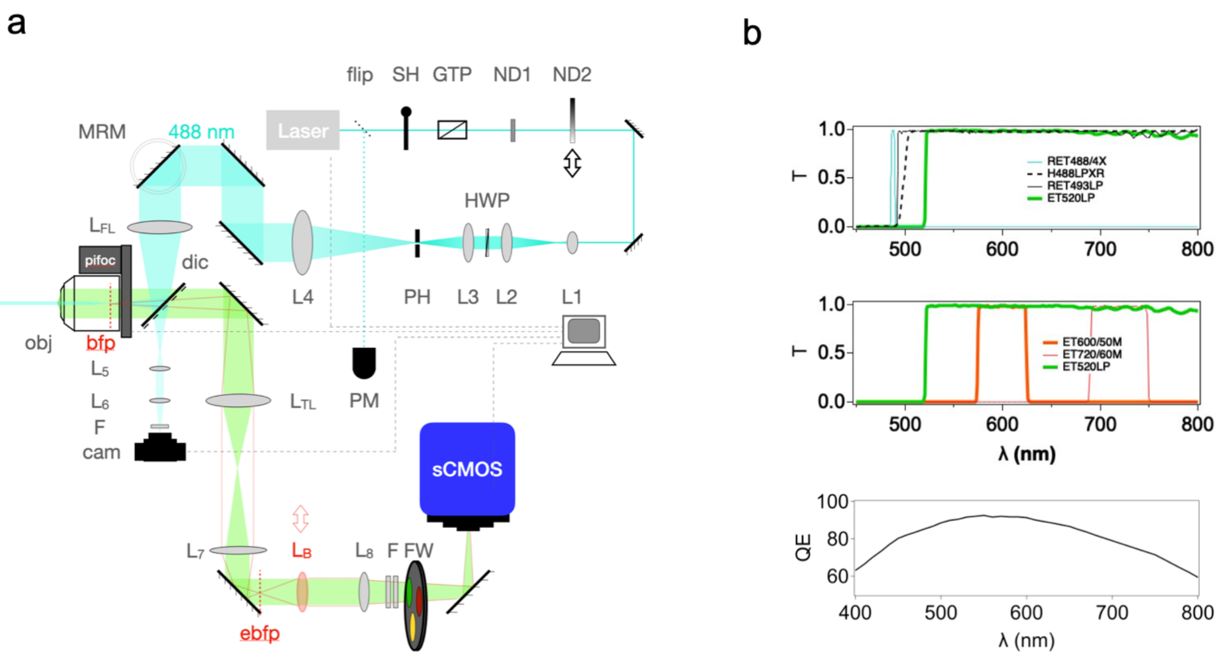


**Fig. S2**. *Combined TIR-SAF microscope, filter and detector spectra.* (**a**), simplified layout of the custom inverted microscope assembled from optical-bench components, see **Supporting experimental procedures** for details. Abbreviations: flip - flippable mirror, SH - shutter, GTP - Glan-Thompson polarizer, ND - neutral-density filter, L - lens, PH - pinhole, MRM - mechanical rotation mount, $L_{FL}$ - focusing lens, dic - dichroic mirror, pifoc - piezo-electric focus drive, obj - microscope objective, $L_{TL}$ - tube lens, (e)bfp - (equivalent) back-focal plane, $L_B$ - motorized Bertrand lens, F - filters, FW - motorized filter wheel, sCMOS - scientific complementary metal-oxide sensor (detector). turquoise and green shading indicate, respectively, excitation and emission optical paths. Red dash shows effect of phase telescope when the Bertrand lens is in place. (**b**), optical characteristics of the key optical components. *Top,* filter spectra showing transmitted intensity (T) vs. wavelength for the laser clean-up, primary dichroic and two long-pass (LP) emission filters always in place (marked "F" on panel a) to reject the back reflected laser beam. *Middle*, additional band-pass (BP) emission filters were housed in a motorized filter wheel (FW), and allowed, respectively selecting for the yellow-green, orange and deep-red spectral bands, respectively. *Bottom*, quantum efficiency of the PCO.edge4.2 sCMOS detector, chip and entry window combined (manufacturer data).

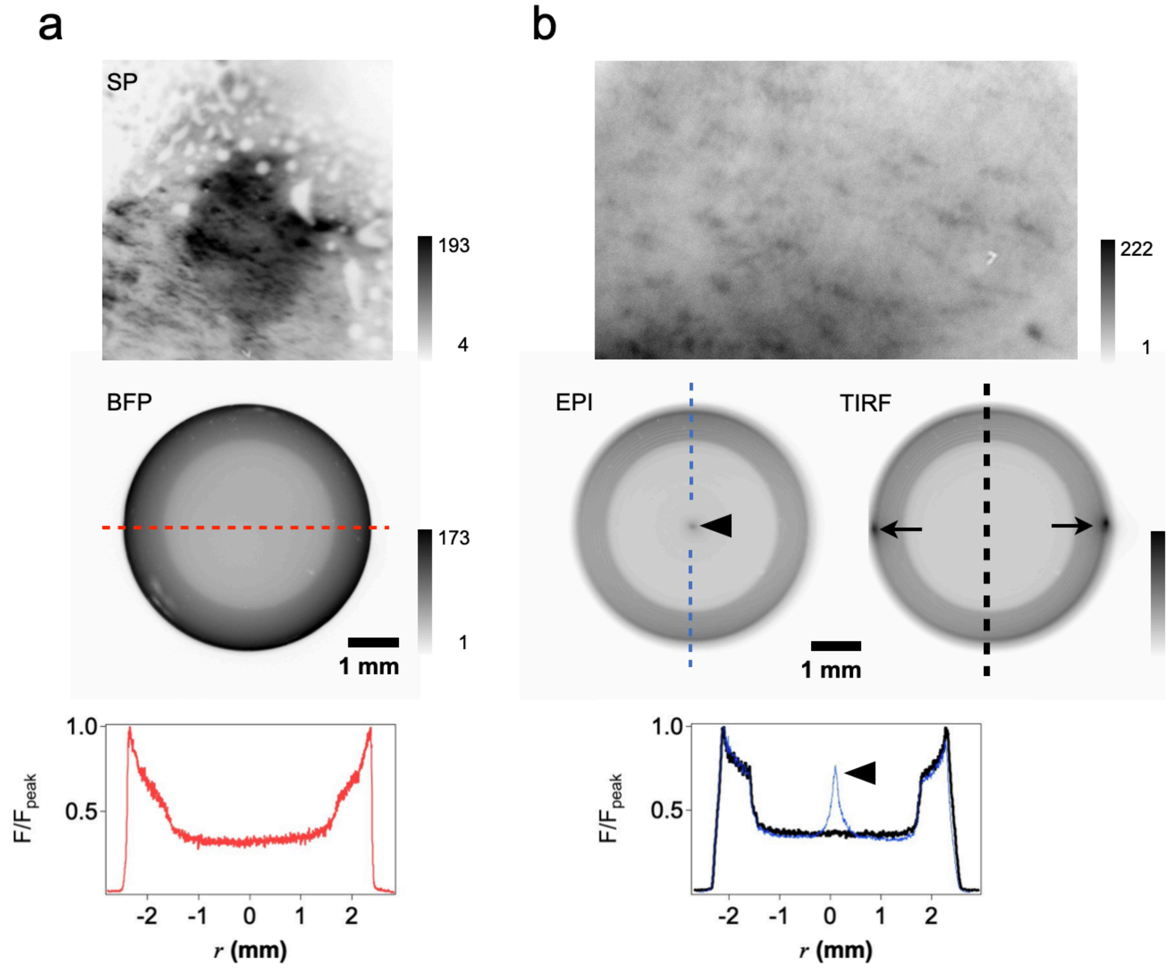


**Fig. S3**. *Routine alignment and measurement steps prior to experiments, for epifluorescence and evanescent-wave fluorescence excitation.* (**a**), 'Quick & dirty' measurement with a text marker (pyranine) smear on a BK-7 microscope coverslip were performed before each experiment to verify, (*i*), the alignment of the excitation path; (*ii*), the symmetry of the of the emission pattern and, (*iii*), a non-obstructed NA and functioning of the motorized Bertrand lens. *Top*, sample-plane (SP) and corresponding back-focal plane (BFP) image, *middle* of a pyranine dye layer on the same type of #1.5 BK-7 glass coverslips as used for the polymer samples, in air. Exposure time ($t_{exp}$) was 500 ms, laser power 4 µW (ND 1), emission filters were a RET493LP and ET520LP with no extra band-pass filters. *Bottom*, normalized intensity profile along the cross-sectional line shown in *red dash* on the BFP image. Image contrast inverted for better clarity. (**b**), *top*, SP image of a TPPS-layer deposited on top of a 65-nm thick My-133-MC Polymer spacer upon epifluorescence excitation (EPI, *middle left*) or total internal fluorescence (TIRF, *right*) excitation. Note the tiny intensity spots at locations corresponding to the focused laser beam (solid arrowhead at ($\vartheta$, $\varphi$)= (0, 0) on EPI image and arrows for the incoming and reflected laser beam on the TIRF image, respectively. $t_{exp}$ = 200 ms, $P$ = 40 µM, filters as above with an additional ET720/60BP filter. *Bottom*, vertical cross-sections (to avoid artefacts from the laser spots) along the lines shown in blue and black, respectively. Note the sharper intensity transition compared to (a), due to the thinner dye layer and more defined emission-wavelength range. As expected, the radiation pattern was independent irrespective of whether fluorescence was excited in EPI or TIRF.

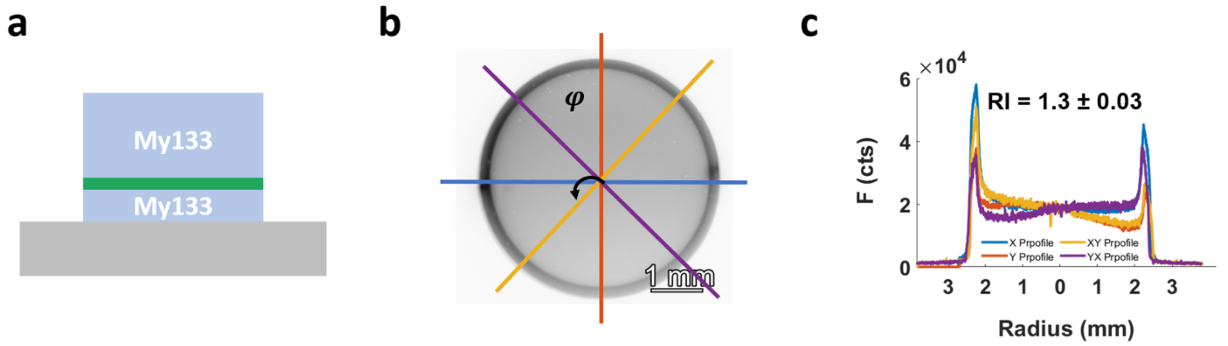


**Fig. S4**. *Emission critical angles measured from intensity cross-sections of BFP images were robust against coverslip tip-tilt.* (a), schematic representation of the studied spacer-dye-sandwich, My-133-MC ( MyPolymer) $n_1$ = 1.33, moisture cured. *Green line* is a 5-nm thick $H_6$-TPPS J-aggregate emitter layer. (b), corresponding BFP image (ex/em = 488 nm/520 nm, LP, contrast inverted for better clarity). Azimuthal orientation of the cross-sections is color coded for line profiles measured at $\varphi$ = 0°, 45°, 90° and 135°, respectively. Pixel sizes were scaled for the two oblique profiles by a factor of $(2)^{½}$ compared to the ones at 0° and 90° for distance measurements. Although slightly different in their intensity, the resulting intensity line profiles, shown in (c), had identical peak positions, critical angles and limiting NAs, even in the presence of small coverslip tip-tilt with respect to the optical axis. In all cases, the measured refractive index (RI) was close to 1.3 within 0.03 RI units.

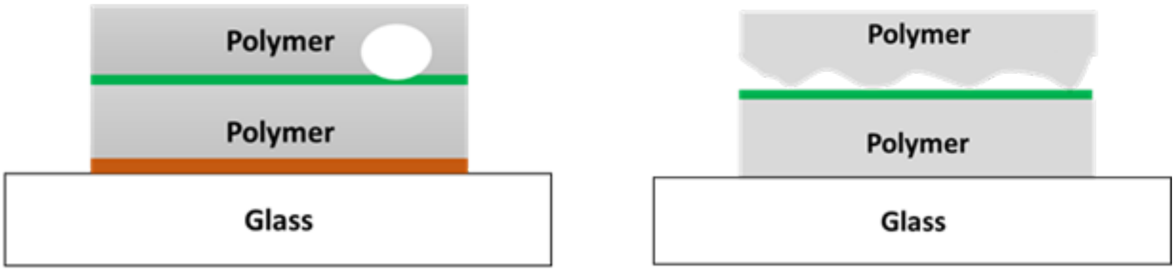


**Fig. S5**. *Possible explanations for a double-ring emission pattern observed in the BFP.* Schematic illustrations show air bubble in the spacer layer (*left*) or an imperfect coverage of the TPPS-dye layer (*right*) as possible nanofabrication defaults in the My-133-MC polymer layer. Drawings are not to scale.

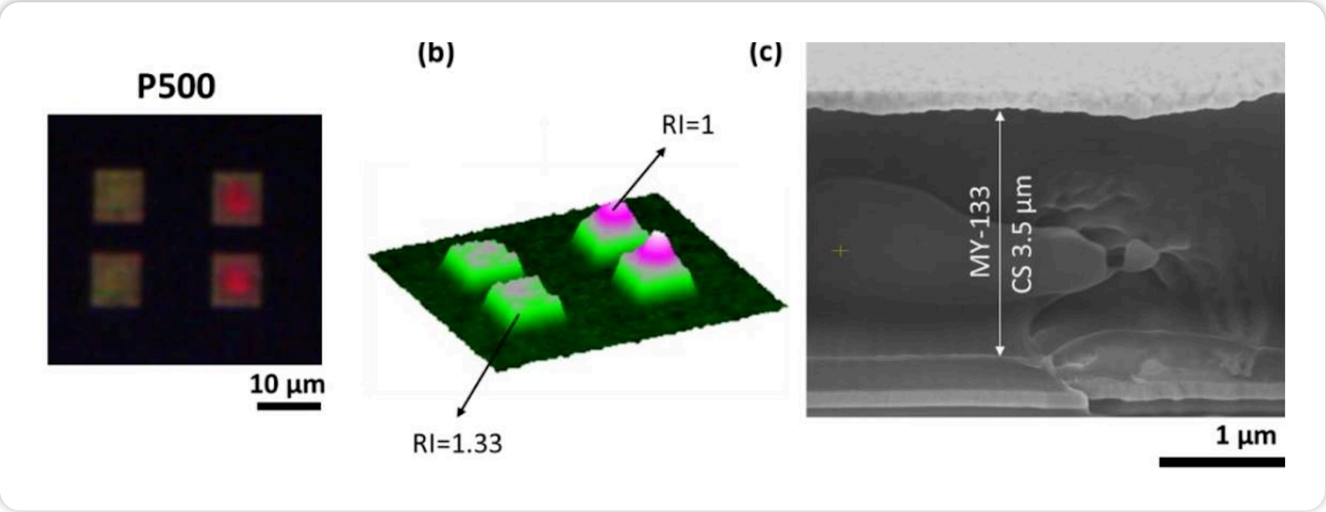


**a** **b** **c**

**Fig. S6**. *Dual-refractive index environments were also detected with nanoplasmonic analysis methods.* (**a**), true-color white-light transmitted image of a plasmonic hole array similar to that shown in fig.(3b) in the main text, with 500-nm periodicity. A 200-nm silver film was deposited on the BK-7 coverglass substrate and covered with a thick (µm-) layer of My-133-MC polymer. Structural colors reflect the plasmonic modes of excitation, related to the geometrical parameters of the plasmonic structure and the RI of the surrounding medium. As the RI was not homogenous for this polymer coverage, different colors are seen on the left and right for the same periodicity of the hole array. (**b**), same in pseudo-color intensity heat map. (**c**), Electron micrograph cross section of the measured sample confirming the presence of cavities within the deposited polymer of this particular layer.

#### SUPPLEMENTARY EXPERIMENTAL PROCEDURES

*Ethics statement.* This study used 2 adult female C57/BL/6 mice (Charles River) and 6 mouse pubs. Mice were bred and housed in the local animal facility on an inverted light-day cycle and with *ad libitum* access to food and water. All experimental procedures were performed in accordance with the French legislation and in compliance with the European Community Council Directive of November 24, 1986 (86/609/EEC) for the care and use of laboratory animals. The used protocols were approved by the local ethics committee.

*Cell preparation and solutions.* Astrocyte cultures were prepared from postnatal-day zero (P0) to P1 mouse cortex as described earlier (Li PNAS 2008, +++REF is the same as in the main text https://www.pnas.org/doi/10.1073/pnas.0909109106). Briefly, cerebral cortices were dissected, the meninges removed and the tissue mechanically dissociated. The obtained cells were filtered, resuspended and plated in plastic dishes and cultured for 1 week at (37°C 5% $CO_2$, 95%) to reach confluence. The then dense cultures were trypsinized, resuspended and transferred onto BK-7 coverslips at 10,000 cells/ml. We used the same bare #1.5 BK-7 glass, like for the polymer samples, with no further coating or treatment. Isolated astrocytes or cells in small islands were recorded during day 2 to 6 after transfer into secondary culture. Recordings were made on a home-built upright microscope (see below) for overviews and on the TIR-SAF optical-bnch microscope (Fig.S2) at RT (20-22°C) under constant perfusion at 0.5–1 ml/min with physiological saline containing (in mM): 140 NaCl, 5.5 KCl, 1.8 $CaCl_2$, 1 $MgCl_2$, 20 glucose, 10 HEPES (a pH buffer). pH was adjusted to 7.3 by the drop-by-frop addition of NaOH. Perfusion was stopped during incubations to label cells with FM dyes.

*Fluorescent markers.* We labeled cultured mouse cortical astrocytes with the FM1-43® (FM3-23®) analog Synaptogreen C4 (C18, Biotium, Fremont, CA) by incubating cells in a static bath with 6.7 µM dye during 2-5 min. FM staining of external plasma membranes was washed off by thoroughly rinsing the cells during 10 min before viewing. *Nota bene* : unlike in neurons, FM labels different subcellular organelles, has a different entry mechanism and a residual cytosolic location in cortical astrocytes. Refer to +++REF Li PNAS 2008 for details.

*Epifluorescence microscopy.* Overview images of FM-labelled astrocytes shown in Fig. 6a and Fig. S7 were taken on a home-built upright microscope described elsewhere +++REF van 't Hoff, Reuter PCCP 2009 https://pubs.rsc.org/en/content/articlelanding/2009/cp/b823155a/unauth. Briefly, the output of a Polychrome II light-source (TILL Photonics, Gräfelfing, Germany) was directed via a fused-silica fiber (NA 0.22) to a custom illuminator, and the fiber exit was imaged into the sample plane (critical illumination). A dual-band GFP-DsRed HC filter set (AHF F56-420) was used, and the excitation wavelength set to 488 nm. Fluorescence was collected through the same ×63/NA0.95w Ph3 dipping lens (Achroplan, ZEISS, Oberkochen, Germany) and imaged via a 180-mm tube lens (Olympus, Hamburg, Germany) and ×2 after-magnification onto the chip of an EMCCD camera (QuantEM 512SC, Photometics, Tucson, AZ). The effective pixel size in the sample plane was 116.5 nm/px. Images were acquired at the highest hardware gain (G=3) and 900 EM-gain, with 100-ms exposure time and 10 MHz readout rate. Unless otherwise stated, time series of 180 images were taken at 1 Hz, and 100 µM ATP were applied to the bath at 1 min to raise the intracellular calcium via purinergic-receptor activation and to test cell viability. Images were subtracted with an average dark image (100 frames taken with the same settings but the shutter closed) to correct for spurious room light and detector intensity offset resulting from dark and amplification noise. All microscope components were controlled with METAMORH software (Molecular Devices, San José, CA). Once a standard recording protocol was established from these overview experiments (1 Hz, rest, 100 µM ATP stimulation, wash), cells were imaged on our TIR-SAF setup (see below).

*TIR-SAF fluorescence microscopy*. A combined TIR-SAF setup was assembled from optical bench components similar to what has been described earlier +++REFs van 't Hoff OE 2008 : https://opg.optica.org/oe/fulltext.cfm?uri=oe-16-22-18495&id=173041; Brunstein BJ 2014a: https://www.sciencedirect.com/science/article/pii/S0006349514000794 , see **Fig. S2a**. Briefly, the beam of a 488-nm beam was expanded and focused in the BFP of an aplanatic and achromatic high-numerical aperture oil-immersion objective (α Plan-Apochromat ×100/NA1.46 oil DIC M27, ZEISS, Oberkochen, Germany). A scanning mirror in a conjugate pupil plane of the excitation optical path permitted adjusting the beam angle and switch between 0° (epifluorescence, EPI) and ~75° (the exact value is limited by the laser spot size and the aperture of the objective). The effective numerical aperture of the objective was measured as $NA_{eff}$ = 1.465± 0.009 +++REF Dai Enderlein OL: https://opg.optica.org/oe/abstract.cfm?uri=OE-13-23-9409, Brunstein BJ 2014a: SAME AS ABOVE. Fluorescence was extracted by 2-mm thick 488-nm super-flat long-pass dichroic mirror (AHF, Tübingen, Germany) was imaged onto a back-illuminated sCMOS detector (Edge4.2 bi, PCO, Kelheim, Germany). A flippable Bertrand lens permitted toggling between sample-plane (SP) and back focal plane (BFP) acquisitions. All steering mirrors were 1" or 2" broadband- (E02-) coated metal mirrors (Thorlabs). The primary dichroic was held in a custom mirror fixing device for Microbench (AHF, F92-325). The residual 0.5%-transmission of the dichroic mirror in continuity of the excitation optical path was filtered (FB488/10M) imaged onto a small DCC1545 CMOS camera for excitation BFP analysis (L5, L6, F, CAM on **Fig. S2**).

*BFP image analysis*. For the aplanatic objective lens used in this study, Abbe's sine condition relates radial distances $r$ measured in the BFP to angles $\vartheta$ via $r=f.n_2\sin(\vartheta_2)$. Here, $f$ and $n_2$ are, respectively, the objective lens' focal length (1.65 mm) and the substrate RI (1.5125). $\vartheta_2$ is the polar beam angle in medium $n_1$, i.e., inside the front lens of the objective or the glass of the substrate. This expression reduces to $r_c=f.n_1$ at the emission critical angle $\vartheta_c$ (Snell), and it becomes $r_{max}=f.NA_{eff}$ at the limiting effective NA. We used the curvature zeros of an equatorial intensity line profile as a measure of $r_c$, related to the local RI ($n_1=r_c/f$, see +++REF Brunstein 2017: https://pubmed.ncbi.nlm.nih.gov/28494964/).

Alternatively, an area-detection algorithm was implemented in MATLAB (The Mathworks, Natick, MA), which first delineated the three BFP-image zones: background, super- and under critical emission components by multiple thresholding. It thus provides average background intensity, and, respectively, the cumulative intensities (after background subtraction) of the SAF and UAF image zones, as well as their ratio, $R = I_{SAF}/I_{UAF}$. A first $R$ estimate is generated using the calculated equivalent radius of the circular areas limited by the emission critical angle, and an *a priori* knowledge about the (measured) effective NA of the objective, $NA_{eff}$ and the previously determined RI of My-133-MC polymer. Error bars were generated by repeating this integration in the range RI +/- 0.05 RI units.
